# Comparison of Osteoblast Calcification in Bio-Oss, Cerasorb, Pro Osteon, and Bio-Tiss Cerabone

**DOI:** 10.64898/2026.05.12.724627

**Authors:** Avin Ghasem, Shirin Zahra Farhad, Maryam Ostadsharif

**Author notes:** Correspondence to: Department of Periodontology, Faculty of Dentistry, Isfahan (Khorasgan)Branch, Islamic Azad University, Isfahan, Iran.

## Abstract

**Background:** Bone graft biomaterials play a critical role in bone regeneration by influencing osteoblast differentiation and mineralization. However, comparative data regarding the osteogenic potential of commonly used graft materials under standardized conditions remain limited.

**Method and material:** In this in vitro experimental study, osteoblast-like cells (MG-63) were cultured with four bone graft materials, including Bio-Oss, Cerasorb, Bio-Tiss Cerabone, and Pro Osteon. The relative mRNA expression of osteogenic markers (COL1 and OPN) was evaluated at 1, 7, 14, and 21 days using real-time PCR. Alkaline phosphatase (ALP) activity and mineralization capacity were also assessed using colorimetric assay and Alizarin Red staining. Data were analyzed using one-way ANOVA and Tukey post hoc test (P < 0.05).

**Results:** Significant differences were observed among the tested materials across all evaluated parameters. Bio-Oss and Cerasorb demonstrated higher gene expression levels and ALP activity compared to Bio-Tiss Cerabone and Pro Osteon (P < 0.05). Mineralization analysis showed significantly greater calcium deposition in the Bio-Oss and Cerasorb groups, whereas Pro Osteon consistently exhibited the lowest osteogenic performance.

**Conclusion:** Bone graft biomaterials significantly influence osteogenic activity in osteoblast-like cells. Bio-Oss and Cerasorb showed superior osteogenic potential, while Pro Osteon demonstrated weaker performance. These findings highlight the importance of material properties in optimizing bone regeneration.

## Introduction

Bone regeneration remains a major challenge in modern dentistry, particularly in implantology and periodontal therapy, where adequate bone volume and quality are essential for successful treatment outcomes (1,15). Following tooth loss, rapid alveolar bone resorption occurs, with studies reporting up to 50% reduction in bone volume within the first year, significantly compromising implant placement and long-term stability (2). Therefore, the use of bone graft materials has become a fundamental strategy in regenerative procedures.

Bone graft substitutes serve as scaffolds that facilitate osteoblast adhesion, proliferation, and differentiation, ultimately promoting new bone formation (3,8). A wide range of graft materials, including xenogeneic, synthetic, and coral-derived biomaterials, have been developed, each with distinct physicochemical properties that influence their biological performance (4,15). Xenografts such as Bio-Oss and Bio-Tiss Cerabone are widely used due to their structural similarity to natural bone and their ability to maintain volume over time (5). In contrast, synthetic materials like Cerasorb (β-tricalcium phosphate) promote osteogenesis through controlled resorption and ion release (6,10,12). Coral-derived materials such as Pro Osteon exhibit a porous architecture resembling cancellous bone; however, concerns remain regarding their resorption behavior and long-term stability (7,14).

Despite extensive research on bone graft materials, several limitations persist in the current literature. Most previous studies have evaluated these biomaterials individually or have focused on a single biological parameter (8). However, osteoblast calcification is a complex, multi-stage process that involves early enzymatic activity, extracellular matrix production, and late-stage mineral deposition (9). Therefore, relying on a single marker may not provide a comprehensive understanding of the osteogenic potential of these materials.

In addition, direct comparative studies evaluating multiple clinically relevant graft materials under standardized experimental conditions remain limited (8). This gap reduces the ability to make evidence-based decisions in selecting the most appropriate graft material for clinical applications.

Accordingly, the novelty of the present study lies in providing a comprehensive, multi-parameter comparison of four widely used bone graft materials (Bio-Oss, Cerasorb, Pro Osteon, and Bio-Tiss Cerabone) within a controlled in vitro model. Osteoblast calcification was evaluated using a combination of biochemical (ALP activity), histochemical (Alizarin Red staining), and molecular (gene expression of COL1 and OPN) markers, allowing for a more integrated assessment of osteogenic behavior.

Therefore, the aim of this study was to compare the osteogenic potential of these materials and to identify the most effective graft in supporting osteoblast calcification under standardized conditions.

## Materials and Methods

This in vitro experimental study evaluated the osteogenic activity of four bone graft biomaterials (Bio-Oss, Cerasorb, Bio-Tiss Cerabone, and Pro Osteon) on osteoblast-like cells over time. All materials were used according to the manufacturers’ instructions.

Human osteoblast-like cells (MG-63 line) with cell code C116 were obtained from the Pasteur Institute of Iran (Tehran, Iran). The cells were cultured in Dulbecco’s Modified Eagle’s Medium (DMEM; high glucose) supplemented with 10% fetal bovine serum and 1% penicillin– streptomycin, and maintained at 37°C in a humidified atmosphere containing 5% CO_2_. Cells at passages 4–8 with viability greater than 90% were used in the study.

Cells were seeded at an appropriate density and exposed to each biomaterial. A total of 35 experimental units (5 groups × 7 replicates) were included. Samples were randomly assigned to five groups using block randomization. Evaluations were performed at 1, 7, 14, and 21 days.(13)

Alkaline phosphatase (ALP) activity was measured at days 7 and 14 using a colorimetric assay kit, and absorbance was recorded at 405 nm.

Mineralization was assessed at day 21 using Alizarin Red S staining (2%, pH 4.2). Calcium deposits were imaged and quantified following dye extraction.

Total RNA was extracted at each time point using a commercial kit. RNA purity and concentration were assessed spectrophotometrically (A260/280). Complementary DNA (cDNA) was synthesized using a reverse transcription kit. Quantitative real-time PCR (qPCR) was performed using SYBR Green chemistry. The expression levels of collagen type I (COL1) and osteopontin (OPN) were measured, with GAPDH as the housekeeping gene. Relative gene expression was calculated using the 2^-ΔΔCt method.(16)

Data were expressed as mean ± standard deviation from at least three independent experiments. Statistical analysis was performed using one-way ANOVA followed by Tukey post hoc test. A P value < 0.05 was considered statistically significant.

## Results

Alkaline phosphatase (ALP) activity increased over time in all experimental groups. At day 7, the highest activity was observed in the Bio-Oss group, followed by Cerasorb and Bio-Tiss Cerabone, while Pro Osteon showed the lowest values. Similar trends were observed at day 14, with Bio-Oss maintaining the highest level and Pro Osteon remaining the lowest. Differences between groups were statistically significant (P < 0.05).

The relative mRNA expression of COL1 showed time-dependent variations among the studied groups (Table 1). At day 1, Bio-Oss showed higher expression levels, while Pro Osteon showed lower levels. At day 7, Bio-Oss and Cerasorb exhibited comparable expression levels and were higher than Bio-Tiss Cerabone and Pro Osteon. At day 14, Cerasorb showed higher expression, whereas Pro Osteon remained lower. At day 21, all groups showed reduced expression levels; however, Bio-Oss and Bio-Tiss Cerabone maintained higher values compared to Pro Osteon. Differences between groups were statistically significant (P < 0.05).

**Table 1.** Mean ± standard deviation of osteoblast cell differentiation based on Collagen I (COL1) gene expression (PCR analysis) in four groups.

| Group | Day 1 | Day 7 | Day 14 | Day 21 |
| --- | --- | --- | --- | --- |
| Pro Osteon | 1.080 ± 0.130 | 2.122 ± 0.276 | 3.584 ± 0.154 | 6.650 ± 0.105 |
| Cerasorb | 1.040 ± 0.027 | 2.996 ± 0.086 | 5.336 ± 0.155 | 7.866 ± 1.287 |
| Bio-Oss | 0.718 ± 0.143 | 3.008 ± 0.082 | 5.284 ± 0.148 | 8.996 ± 0.171 |
| Bio-Tiss Cerabone | 1.000 ± 0.031 | 2.270 ± 0.047 | 4.508 ± 0.138 | 7.874 ± 0.138 |

Group effect: $F = 42.35$ , $P < 0.001$ | Time effect: $F = 1458.91$ , $P < 0.001$ | Interaction effect: $F = 11.43$ , $P < 0.001$

The relative mRNA expression of OPN also demonstrated time-dependent variations (Table 2). At day 1, higher expression levels were observed in the Bio-Oss group, while Pro Osteon showed lower levels. At day 7, Bio-Oss and Cerasorb exhibited higher expression compared to Bio-Tiss Cerabone and Pro Osteon. At day 14, Cerasorb showed the highest expression, whereas Pro Osteon remained the lowest. At day 21, expression levels decreased in all groups, with Bio-Oss and Bio-Tiss Cerabone showed relatively higher values compared to Pro Osteon. Differences between groups were statistically significant (P < 0.05).

**Table 2.** Mean ± standard deviation of osteoblast cell differentiation based on Osteopontin (OPN) gene expression (PCR analysis) in four groups.

| Group | Day 1 | Day 7 | Day 14 | Day 21 |
| --- | --- | --- | --- | --- |
| Pro Osteon | 2.740 ± 0.232 | 2.438 ± 0.150 | 2.232 ± 0.124 | 2.136 ± 0.129 |
| Cerasorb | 3.562 ± 0.091 | 3.722 ± 0.068 | 3.420 ± 0.102 | 2.618 ± 0.085 |
| Bio-Oss | 3.878 ± 0.097 | 3.642 ± 0.077 | 3.216 ± 0.099 | 3.030 ± 0.110 |
| Bio-Tiss Cerabone | 3.152 ± 0.092 | 3.086 ± 0.057 | 3.108 ± 0.091 | 2.890 ± 0.077 |

Group effect: $F = 355.27$ , $P < 0.001$ | Time effect: $F = 136.50$ , $P < 0.001$ | Interaction effect: $F = 17.39$ , $P < 0.001$

Mineralization assessment using Alizarin Red staining further confirmed these findings (Table 3). Quantitative analysis demonstrated that calcium deposition was significantly higher in the Bio-Oss and Cerasorb groups compared to Bio-Tiss Cerabone and Pro Osteon (P < 0.001). Pro Osteon exhibited the lowest mineralization capacity throughout the study period.

**Table 3.**
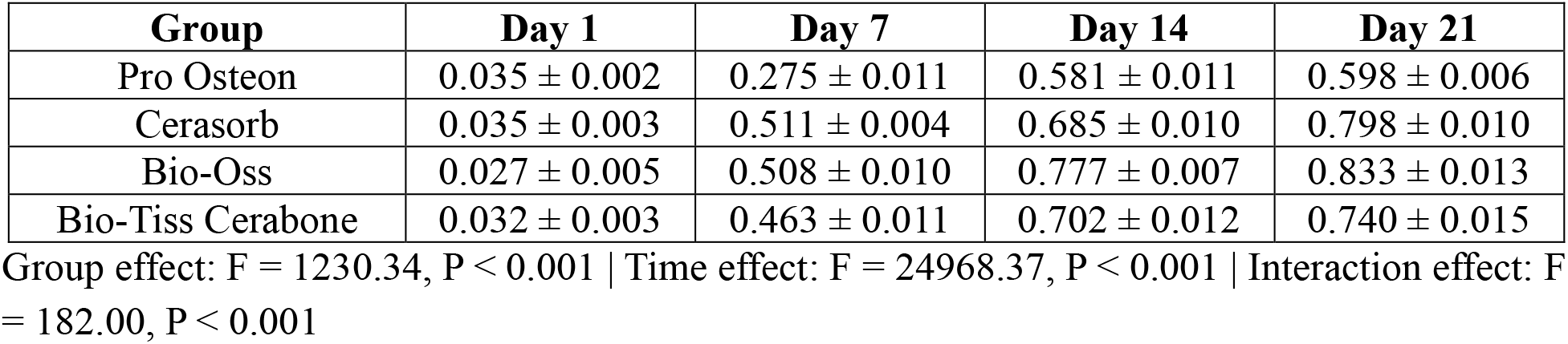
Mean ± standard deviation of osteoblast cell differentiation based on mineralization assay in four groups.

| Group | Day 1 | Day 7 | Day 14 | Day 21 |
| --- | --- | --- | --- | --- |
| Pro Osteon | 0.035 ± 0.002 | 0.275 ± 0.011 | 0.581 ± 0.011 | 0.598 ± 0.006 |
| Cerasorb | 0.035 ± 0.003 | 0.511 ± 0.004 | 0.685 ± 0.010 | 0.798 ± 0.010 |
| Bio-Oss | 0.027 ± 0.005 | 0.508 ± 0.010 | 0.777 ± 0.007 | 0.833 ± 0.013 |
| Bio-Tiss Cerabone | 0.032 ± 0.003 | 0.463 ± 0.011 | 0.702 ± 0.012 | 0.740 ± 0.015 |

Group effect: $F = 1230.34$ , $P < 0.001$ | Time effect: $F = 24968.37$ , $P < 0.001$ | Interaction effect: $F = 182.00$ , $P < 0.001$

Time-dependent analysis indicated a significant effect of time, group, and their interaction on osteogenic outcomes (P < 0.001). While all groups showed an overall increase in mineralization over time, the relative differences between materials remained consistent.

Overall, Bio-Oss and Cerasorb exhibited superior osteogenic performance, Bio-Tiss Cerabone showed moderate activity, and Pro Osteon demonstrated the weakest response among the tested materials.

## Discussion

The findings of the present study demonstrated that the type of bone graft material significantly influences osteogenic activity in osteoblast-like cells. This effect was consistently observed across biochemical (ALP activity), molecular (COL1 and OPN expression), and mineralization outcomes. Among the evaluated biomaterials, Bio-Oss showed the highest osteogenic performance, followed by Cerasorb, whereas Bio-Tiss Cerabone exhibited moderate behavior and Pro Osteon demonstrated the lowest activity.

The superior performance of Bio-Oss can be attributed to its structural and compositional similarity to natural bone, as well as its highly porous architecture, which enhances cell adhesion and proliferation. Recent studies have confirmed that xenogeneic grafts provide a stable scaffold that supports osteoblast differentiation and promotes uniform mineral deposition (5,11). In addition, its low resorption rate allows for prolonged structural support, which is critical for sustained bone regeneration (1,5).

Cerasorb, a β-tricalcium phosphate-based material, exhibited comparable osteogenic potential in several parameters. This finding is consistent with literature highlighting the role of calcium phosphate biomaterials in stimulating osteogenesis through controlled ion release (6,10,12). The release of calcium and phosphate ions has been shown to activate signaling pathways involved in osteoblast differentiation and extracellular matrix mineralization (10,12).

Bio-Tiss Cerabone demonstrated moderate osteogenic activity, which may be explained by differences in its physicochemical characteristics, including porosity, crystallinity, and surface topography. Although it is also a xenograft, variations in processing methods can influence its biological performance. Previous and recent studies have highlighted that microstructural properties of biomaterials play a critical role in regulating cell–material interactions and subsequent bone formation (8,13,15).

In contrast, Pro Osteon showed the lowest osteogenic activity across all evaluated parameters. This may be related to its coral-derived origin and its relatively faster resorption rate, which can compromise scaffold stability over time (7,14). Recent findings suggest that excessive resorption may limit the duration of cellular support and reduce long-term osteogenic efficiency (14).

Overall, the results of the present study are in agreement with previous and recent research indicating that both material composition and structural properties are key determinants of osteogenic potential (3,4,15). The consistent superiority of Bio-Oss and the comparable performance of Cerasorb highlight the importance of scaffold stability, controlled biodegradation, and ion release in optimizing bone regeneration outcomes.

Despite these findings, several limitations should be considered. The in vitro design does not fully replicate the complex in vivo environment, where factors such as vascularization, immune response, and mechanical loading significantly influence bone regeneration. Additionally, only selected osteogenic markers were evaluated, while other molecular pathways involved in bone formation were not assessed. Osteoblast differentiation is a multi-stage process involving early enzymatic activity, extracellular matrix maturation, and late-stage mineral deposition, and evaluating multiple osteogenic markers provides a more comprehensive understanding of cellular responses to biomaterials (9).

Recent studies recommend integrating advanced analytical approaches to better understand biomaterial–cell interactions (13).

Future research should focus on in vivo studies and clinical trials to validate these findings and evaluate long-term performance. Moreover, investigating additional molecular mechanisms and the interaction between biomaterials and the host environment may provide deeper insights into optimizing bone graft materials for clinical applications.

## Conclusion

Within the limitations of this in vitro study, Bio-Oss and Cerasorb demonstrated superior osteogenic performance compared to Bio-Tiss Cerabone and Pro Osteon. The findings highlight the importance of scaffold stability and controlled ion release in enhancing osteoblast activity and mineralization. From a clinical perspective, both xenogeneic and optimized synthetic graft materials may provide favorable conditions for bone regeneration, particularly in procedures requiring predictable osteogenic outcomes. However, further in vivo studies are necessary to confirm these results and support their clinical application.

